# Direct engagement of mycobacterial RadA by c-di-AMP at a non-catalytic site suppresses its DNA-stimulated ATPase activity

**DOI:** 10.64898/2026.10.22.760896

**Authors:** Shreya Shree, Shanavas P Greeshma, Kulhar Nitin, Eerappa Rajakumara

## Abstract

Cyclic di-adenosine monophosphate couples DNA damage sensing to genome maintenance in mycobacteria, but how it regulates homologous recombination machinery beyond RecA remains unclear. Using *Mycobacterium smegmatis* RadA (MsRadA) as a model, we show by isothermal titration calorimetry that c-di-AMP binds to MsRadA, providing biophysical evidence for an interaction previously inferred only from genetic studies. C-di-AMP binding is independent of magnesium ions and is not affected by prior occupancy of the ATP-binding site by the ATP analogue β,γ-methylene adenosine 5′-triphosphate, indicating that c-di-AMP binds at a site distinct from the canonical ATPase pocket. Functionally, c-di-AMP reduces MsRadA’s DNA-dependent ATPase activity. Structural modelling and molecular dynamics (MD) simulations show that c-di-AMP is stably bound in the central channel of the RadA hexamer, non-covalently engaged with the DNA-binding residues Arg197, Lys226, and Arg234, which are contributed by two adjacent subunit interfaces of the hexamer. In the modelled RadA-ATP-c-di-AMP-DNA quaternary complex, c-di-AMP was predicted at a peripheral site near, but distinct from, the ATP-binding pocket. Together, these findings identify RadA as a direct physical target of c-di-AMP within the homologous recombination pathway and provide a framework for testing how c-di-AMP regulates RadA during DNA damage.

## Introduction

Bacteria continuously sense and respond to intracellular and environmental cues through cyclic nucleotide based secondary messengers, including cyclic di-nucleotides (CDN), for example, cyclic di-guanosine monophosphate (c-di-GMP) and cyclic di-adenosine monophosphate (c-di-AMP) are known to integrate multiple external stimuli and eventually transmitting them to different cellular targets. [1–4] Recently, c-di-AMP has emerged as a signaling molecule in pathogenic mycobacteria, where it links fundamental physiological processes – DNA repair metabolism, osmotic homeostasis, potassium homeostasis, and cell wall metabolism – to the cell’s overall physiological state. [5,6] Therefore, structural and functional investigations of c-di-AMP’s macromolecular targets, their effectors as well as the enzymes that regulate cellular concentration of c-di-AMP may be helpful for developing therapeutics to selectively disrupt signaling pathways in pathogens such as mycobacterial species. [5,7]

In *Mycobacterium smegmatis*, c-di-AMP is synthesized by DNA integrity Scanning A (DisA) protein. DisA joins two ATP molecules together using a metal ion to hydrolyze the β and γ phosphates, forming the ring-shaped c-di-AMP and stops doing this whenever it senses damaged DNA. [7,8] It scans the chromosome for branched DNA intermediates formed during replication, repair and recombination. Upon encountering such structures, DisA’s c-di-AMP synthesis activity is inhibited, lowering the cellular level of c-di-AMP. This constitutes a signal of DNA damage detection. [9] Elevated intracellular c-di-AMP also remodels surface-associated behaviour. In several species, loss of c-di-AMP phosphodiesterase activity increases cell-cell aggregation and reduces motility. [10,11] High c-di-AMP promotes biofilm in *Streptococcus mutans.* [12] It also suppresses biofilm formation in *Bacillus subtilis*, and in *M. smegmatis*. [10,13]

The link between c-di-AMP and genome integrity has recently been established through its regulatory role in Recombinase A (RecA) and Radiation sensitive A (RadA) dependent homologous recombination (HR). [14] In *Mycobacterium tuberculosis*, c-di-AMP can bind the recombinase MtRecA and destabilize the nucleoprotein filament intermediate formed during RecA-dependent search and exchange of complementary DNA strands, attenuating downstream SOS signaling. Notably, c-di-AMP attenuates the DNA strand exchange promoted by MsRecA as well as MtRecA through the disassembly of RecA nucleoprotein filaments. [15] RadA and DisA occur in a conserved operon across several bacteria including *Mycobacterium tuberculosis and Mycobacterium smegmatis*, suggesting that RadA functions within the c-di-AMP-mediated DNA damage repair pathway.[16] Bacteria coordinate their HR mediated DNA damage pathways through a network linking RecA, RadA, and DisA with c-di-AMP acting as the regulatory signal that ties them together.[17] The link that connects c-di-AMP and RadA has so far been inferred genetically, but the molecular mechanisms of regulation of RadA by c-di-AMP remain unresolved.[18]

A previous report showed that loss of RadA suppressed the elevated recombination phenotype caused by loss of c-di-AMP synthesis in *Mycobacterium smegmatis*, implicating RadA in a c-di-AMP-regulated pathway. [14] Further genetic comparisons suggested RecA and RadA act within the same pathway. This genetic evidence linked RadA to c-di-AMP signaling. [18] Based on published genetic interaction studies, we hypothesized that c-di-AMP could directly bind to RadA, just like RecA and regulate its activities. Here, we attempt to address this gap by measuring the direct interactions between c-di-AMP and MsRadA using the isothermal titration calorimetry (ITC) binding studies. Following binding studies, we assessed the effect of c-di-AMP on RadA’s catalytic activity using ATPase assays. Further, through structural modeling and molecular dynamics simulation based computational analyses, we mapped the c-di-AMP binding sites in the MsRadA.

Our in vitro findings provide the first biophysical evidence of a direct interaction between MsRadA and c-di-AMP, and biochemically demonstrate that this binding negatively regulates the DNA-dependent ATPase activity of MsRadA. Additionally, structural modeling and MD simulations suggest that binding of c-di-AMP on the subunit interface of the hexameric central channel away from the ATP-binding pocket, involves DNA contact residues Arg197, Lys226 and Arg234, supporting a model in which c-di-AMP regulates RadA’s DNA-dependent ATPase activity allosterically, rather than directly competing at the active site.

## Results

### Magnesium enhances the ATP analogue binding affinity to MsRadA

The DNA-dependent ATPase activity of the recombinase MsRadA depends directly on free magnesium ion (Mg²⁺) concentration alongside the primary ATP-bound cofactor [19]. To investigate whether Mg²⁺ affects the binding of MsRadA, we performed ITC binding studies using recombinant MsRadA and adenosine 5′-(β,γ-methylene) triphosphate (AMP-PCP), a non-hydrolysable ATP analogue (Fig. 1A). The binding affinity of AMP-PCP to MsRadA is 88.9 ± 7.25 µM (Fig. 1B) (Supplementary Table 1). On the contrary, the binding affinity of MsRadA to AMP-PCP in the presence of a physiological concentration of Mg²⁺ is 6.5 ± 1.9 µM (Fig. 1C) (Supplementary Table 1), which is (∼14 fold) more than AMP-PCP without magnesium ions. These results infer that Mg²⁺ is not only required for the DNA-dependent ATPase activity of MsRadA but also enhances the binding affinity of the ATP analogue to MsRadA.

**Figure 1.**
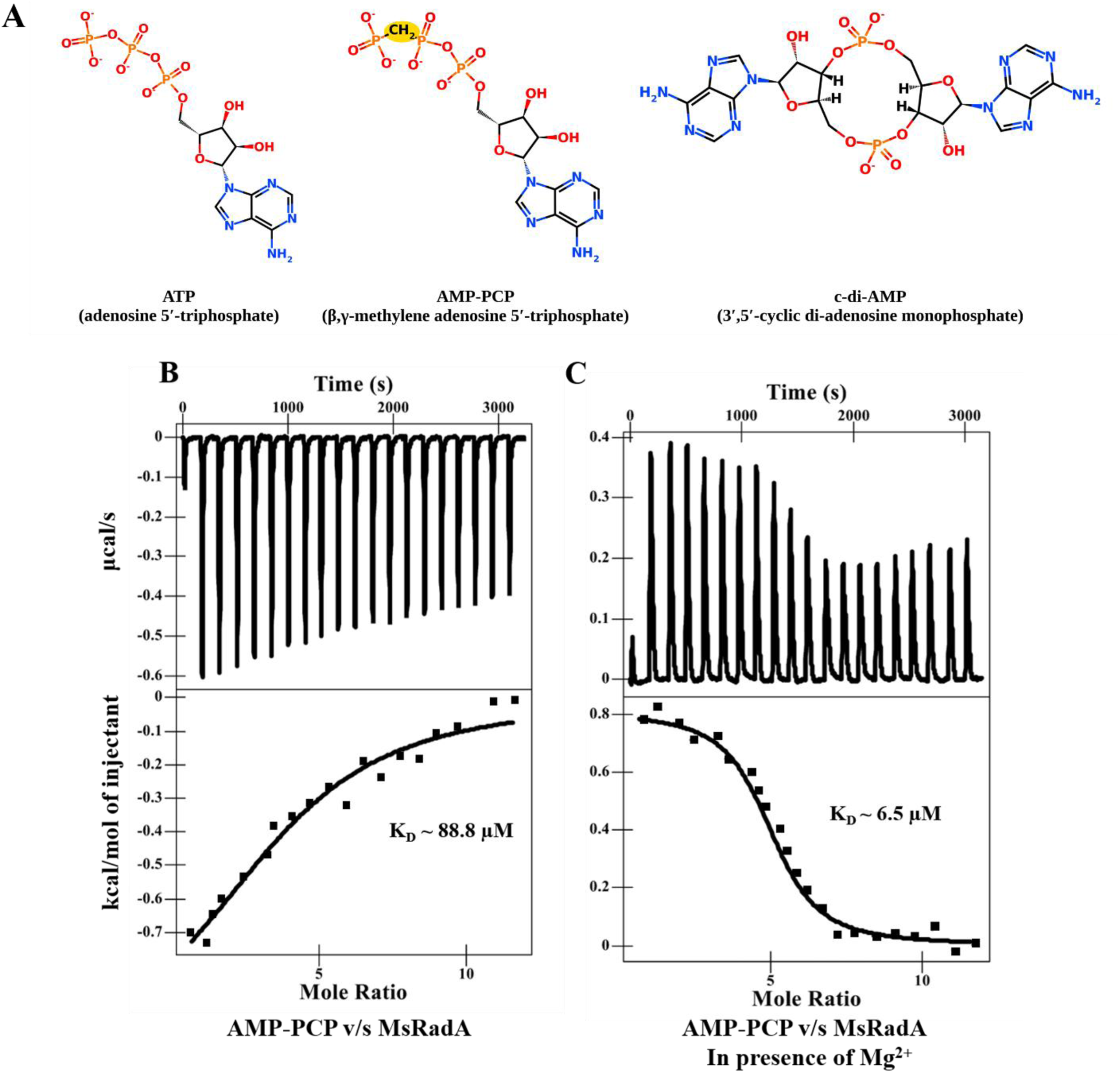
ITC binding studies of an ATP analogue to MsRadA. (A) 2-D structures of the nucleotides: ATP (adenosine 5′-triphosphate), AMP-PCP (β,γ-methylene adenosine 5′-triphosphate) and c-di-AMP (3′,5′-cyclic di-adensosine monophosphate). ITC binding studies of AMP-PCP against MsRadA in the (B) absence and (C) presence of Mg^2+^. Raw ITC data (upper panel) and normalized integration data (lower panel). The measured molar dissociation constant (K_D_) was determined using NanoAnalyze software. Image produced with NanoAnalyze version 3.12.5 (TA instruments).

### C-di-AMP shows direct interaction with MsRadA independent of Mg²⁺

The c-di-AMP binds the RadA partner, the recombinase MtRecA, and destabilises the nucleoprotein filament intermediate formed during RecA-dependent search and exchange of complementary DNA strands. [14,15] Strikingly, the loss of RadA suppressed the elevated recombination phenotype caused by the loss of c-di-AMP synthesis. [18] Based on the above observations, we hypothesized that c-di-AMP directly interacts with RadA to modulate the RecA-RadA-mediated HR activity in *Mycobacteria*. We therefore sought to determine direct physical interactions between c-di-AMP and RadA. ITC binding studies revealed direct interaction with a K_D_ of 14.2 ± 0.6 μM (Fig. 2A) (Supplementary Table 1).

**Figure 2.**
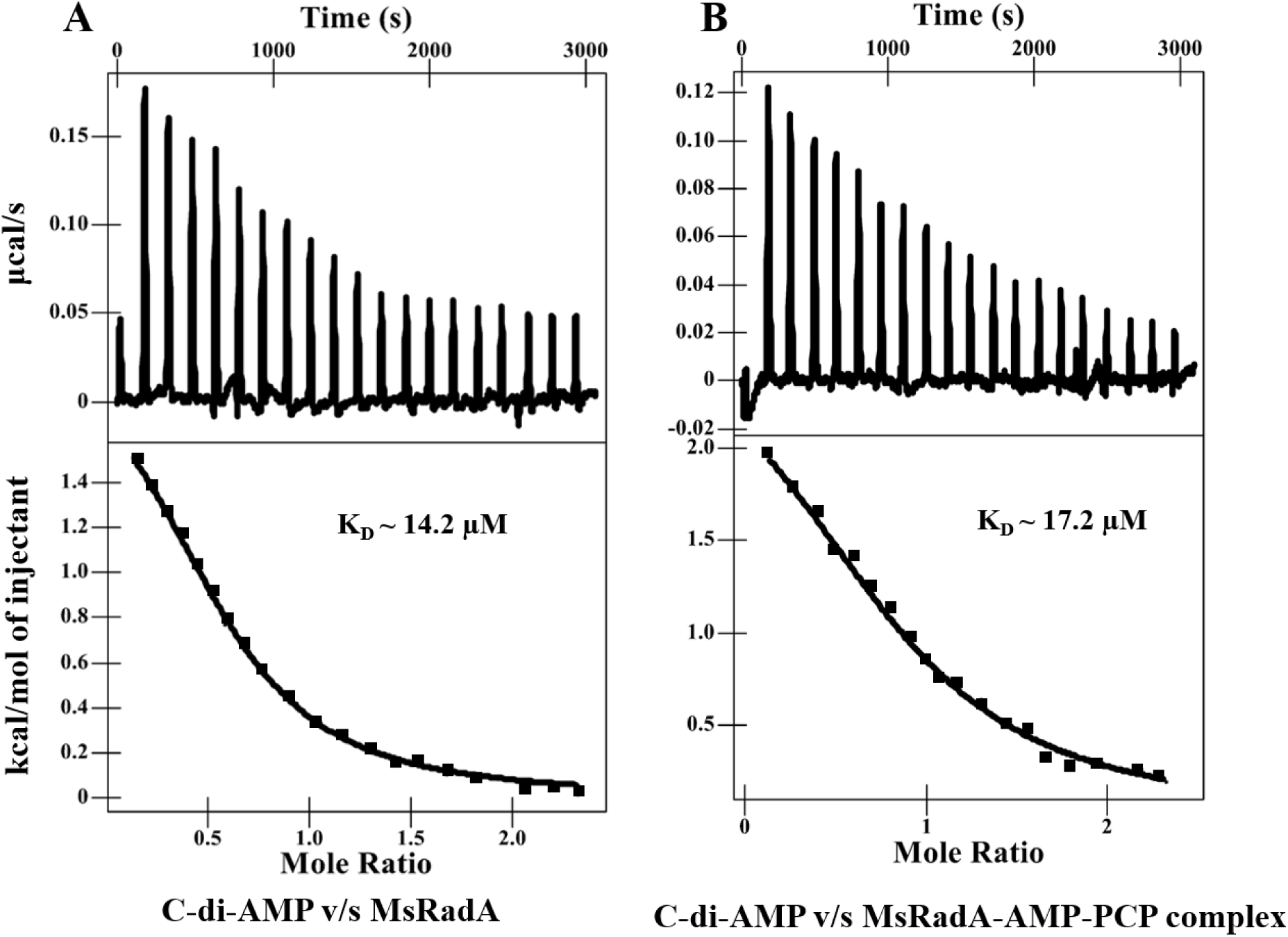
ITC binding studies of (A) C-di-AMP against MsRadA and (B) C-di-AMP against MsRadA-AMP-PCP complex. Raw ITC data (upper panel) and normalized integration data (lower panel). The measured molar dissociation constant (K_D_) was determined using Nano Analyze software. Image produced with NanoAnalyze version 3.12.5 (TA instruments).

As Mg²⁺ is an essential cofactor for nucleotide binding in RecA-family ATPases like RadA, we asked whether it similarly affects binding of the secondary messenger c-di-AMP, we then investigated whether Mg²⁺ influences the c-di-AMP binding activity of MsRadA, like the ATP analog binding of MsRadA. However, there is no significant change in the binding affinity of MsRadA to c-di-AMP in the presence of a physiological concentration of Mg²⁺ (K_D_ ∼ 14.7±0.7 μM), suggesting that c-di-AMP binding to MsRadA is independent of Mg²⁺ (Supplementary Fig. S1). Mg²⁺ coordinates the phosphates of ATP at the canonical ATPase active site, which likely accounts for the higher affinity of MsRadA for the ATP analogue in its presence (Fig. 1C). In contrast, c-di-AMP binding does not show the same Mg²⁺ dependence. This difference in metal requirement suggests that the ATP analogue and c-di-AMP bind to distinct sites on MsRadA.

### AMP-PCP does not compete with c-di-AMP for binding to MsRadA

To directly test whether c-di-AMP and AMP-PCP compete for the same site on RadA, we titrated c-di-AMP against the preformed RadA-AMP-PCP (1:20 molar ratio) complex. C-di-AMP bound the RadA-AMP-PCP complex with a K_D_ ∼ 17 µM (Fig. 2B), closely comparable in both affinity and thermodynamic signature to c-di-AMP binding apo RadA (K_D_ ∼14.2 µM) (Fig. 2A) (Supplementary Table 1). Prior occupancy of the ATP-binding site by AMP-PCP, therefore, has no significant effect on subsequent c-di-AMP binding (Fig. 2A & B).

### C-di-AMP negatively regulates the DNA-stimulated ATPase activity of MsRadA

Wild-type MsRadA did not show ATPase activity in the absence of DNA and was stimulated in the presence of single-stranded DNA, consistent with RadA’s DNA-dependent ATPase mechanism [20]. To determine whether c-di-AMP regulates the DNA-dependent ATPase activity of MsRadA, we measured it in the presence and absence of a fixed concentration of c-di-AMP. In the absence of c-di-AMP, MsRadA exhibited an ATPase activity of ∼10.5±0.62 µM/min (n=3), whereas addition of 0.5 mM c-di-AMP reduced ATPase activity to ∼7.6±0.26 µM/min (n=3), corresponding to an approximately 28% decrease in activity (unpaired t-test, p < 0.01) (Fig. 3). This suggests that c-di-AMP can negatively regulate MsRadA’s catalytic activity.

**Figure 3.**
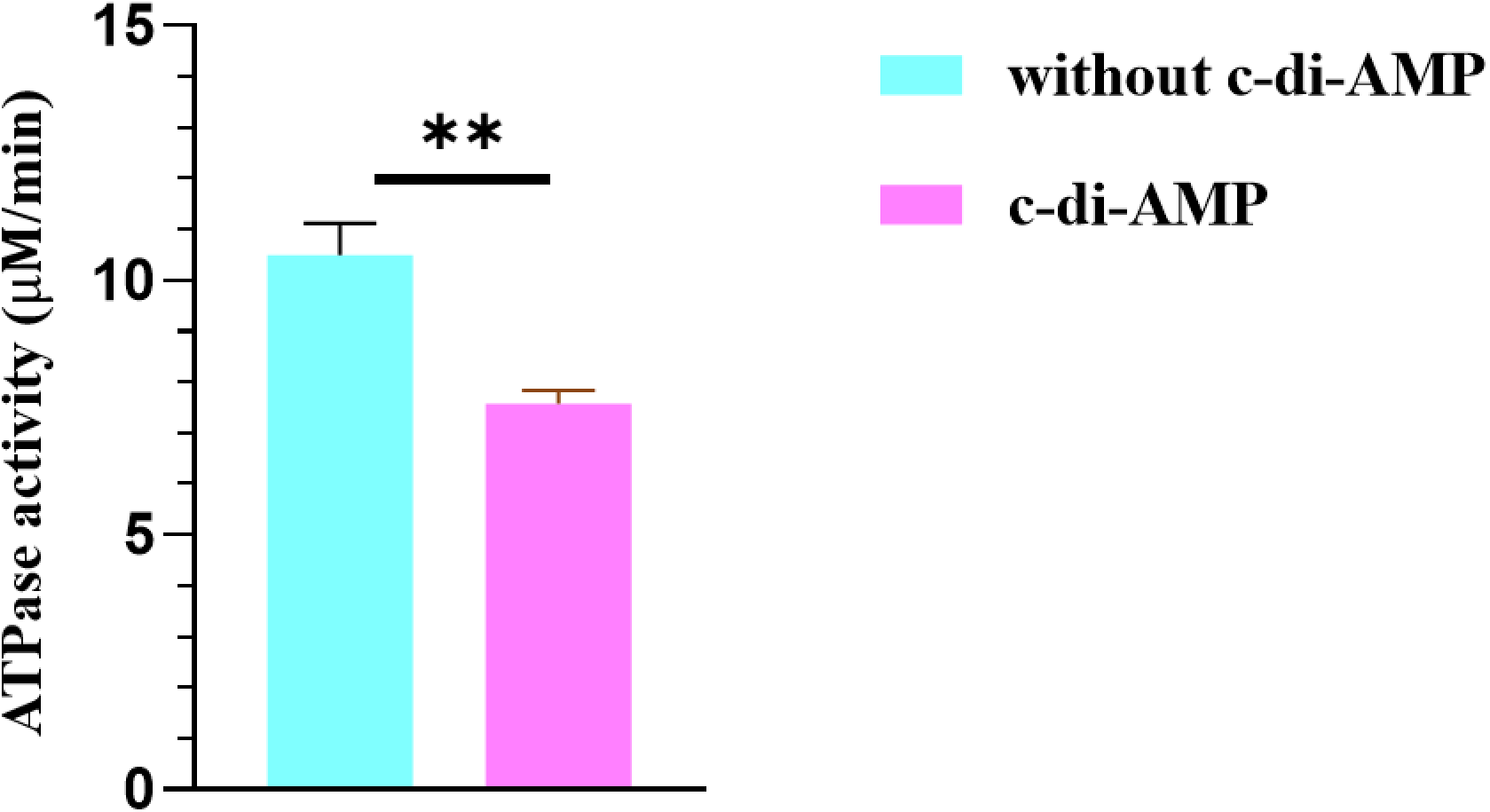
DNA-dependent ATPase activity of MsRadA in presence of c-di-AMP. ATPase activity of MsRadA measured by an NADH-coupled assay in the absence and presence of c-di-AMP. Data are the mean ± SD of n = 3 independent experiments. **p < 0.01, unpaired two-tailed t-test. Data were analyzed and plotted using GraphPad Prism 8.

### C-di-AMP binding site on MsRadA overlaps with the DNA-binding site, distinct from the ATP-binding pocket

Based on ITC binding studies and ATPase assay results, we performed computational studies to determine the c-di-AMP binding site on MsRadA and its relative position compared with ATP and DNA binding sites. We first generated structural models of the MsRadA hexamer using the Protenix server, then of the ternary complexes RadA-ATP-DNA and RadA-ATP-c-di-AMP, and of the quaternary complex RadA-ATP-c-di-AMP-DNA. A duplex DNA with an overhang, which is a substrate for RadA (Supplementary Table 2), was used to model the above RadA complexes [21]. The modeled complexes showed pTM scores of 0.70-0.80 and ipTM scores of 0.66-0.78 ( Supplementary Table 3), indicating moderate confidence in the overall assembly and the interface. [22,23]

In the RadA-ATP-c-di-AMP ternary complex, ATP and c-di-AMP were found to occupy separate positions on every subunit interface (Fig. 4A). This holds true for all six interfaces in the hexamer. In the RadA-ATP-DNA complex, DNA was found to thread through the central channel of the hexamer (Fig. 4B), as found in the cryo-EM structure of *Streptococcus pneumoniae* RadA (SpRadA) with gapped DNA [21]. ATP occupied the expected active site, contacting the Walker A P-loop and an arginine finger from the neighbouring subunit in both models, and Mg²⁺ coordination with ATP and residues from the Walker motif and P-loop is highly similar to reported RadA and RecA structures (Supplementary Fig. S2), suggesting that the modelled complexes represent physiological assemblies. However, c-di-AMP binds at the subunit interfaces of hexameric RadA, about 10-12 Å from the ATP-binding site and close to the central channel.

**Figure 4.**
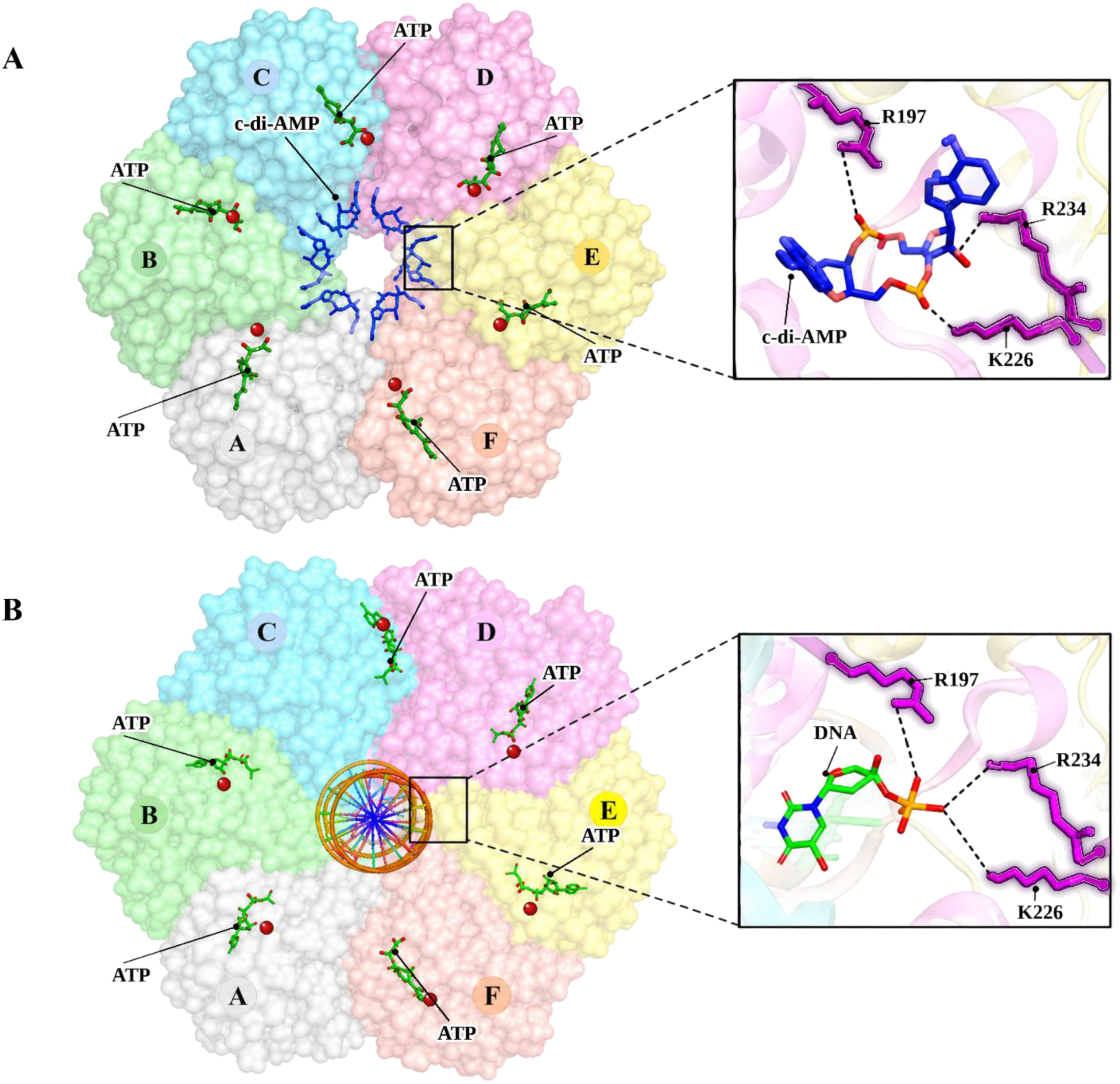
Modelled structures of the MsRadA-c-di-AMP and MsRadA-DNA complexes. (A) Surface representation of the hexamer viewed down the central axis, with protomers A to F colored individually. The ATP bound in each protomer is shown as green sticks, the red sphere is Mg^2+^, C-di-AMP bound in central channel is shown in blue sticks. The boxed region is enlarged in the inset, which shows the phosphate backbone contacted by Arg197, Lys226 and Arg234 (magenta sticks). Black dotted lines mark interactions. (B) DNA modelled in the central channel. In the inset, Arg197, Lys226 and Arg234 (magenta sticks) contact the phosphate groups of the DNA nucleotide. Images were produced using Pymol (The PyMOL Molecular Graphics System).

We analyzed both models using the PLIP server to identify interacting residues. In the c-di-AMP-bound model, the positively charged residues Arg197, Lys226, and Arg234 were found to interact with the phosphate groups of c-di-AMP. In the DNA-bound model, the same positively charged residues, Arg197, Lys226, and Arg234, contact the negatively charged phosphate groups of the DNA (Fig. 4A & B). These residues were common to both models, indicating some overlap between the c-di-AMP and DNA-binding regions within the inner ring of the MsRadA hexamer.

We modelled the RadA-ATP-c-di-AMP-DNA quaternary complex (Supplementary Table 3) to investigate additional c-di-AMP binding sites on RadA besides the DNA binding site. In the complex model (Fig. 5), DNA occupied the central channel as in the RadA-DNA complex (Fig 4B), whereas c-di-AMP was bound at a peripheral site near, but distinct from, the ATP-binding pocket (Fig. 5), unlike the RadA–ATP–c-di-AMP complex (Fig. 4A).

**Figure 5.**
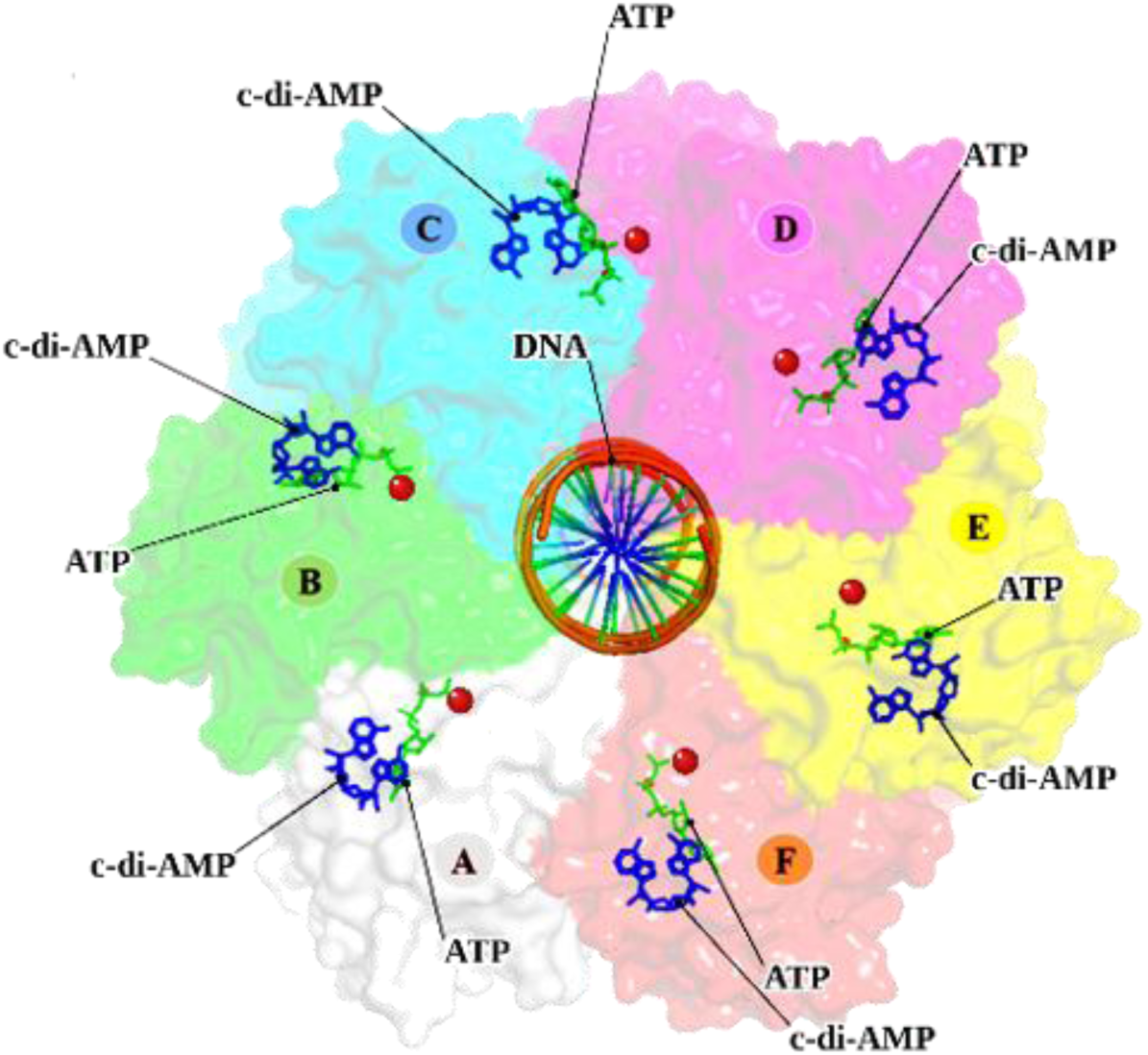
Modelled strcuture of the MsRadA-ATP-c-di-AMP-DNA quaternary complex. MsRadA hexameric ring (subunits A-F, individually colored) modeled with dsDNA with overhang threaded through the central channel. Green sticks at each subunit interface represent ATP occupying the canonical nucleotide-binding pocket. C-di-AMP (blue sticks) is at the periphery near the ATP binding pocket in this model. Images were produced using Pymol (The PyMOL Molecular Graphics System).

### Molecular dynamics suggests c-di-AMP does not perturb ATP positioning in the nucleotide-binding site

To examine how c-di-AMP and ATP occupy the RadA hexamer, we performed 250 ns all-atom MD simulations of the MsRadA-ATP and MsRadA-c-di-AMP binary complexes and MsRadA-ATP-c-di-AMP ternary complex (Supplementary Table 3). The protein RMSD plateaued at ∼4-5 Å across all three systems after 100 ns (Fig. 6A & B), indicating comparable global stability regardless of ligand occupancy. Ligand RMSD was analyzed at the C–D interface. ATP RMSD was ∼4–5 Å both in the presence and absence of c-di-AMP (Fig. 6A), indicating that c-di-AMP does not substantially alter ATP positioning at the Walker A site. In the simulations, c-di-AMP remained bound at the subunit interface in both conditions, although its RMSD was higher with ATP (∼7 Å) than without (∼2-3 Å) (Fig. 6B). This suggests that ATP occupancy of neighboring sites changes the flexibility of the bound c-di-AMP. Together, these simulations are consistent with a model in which c-di-AMP does not act by direct competition at the ATP pocket.

**Figure 6.**
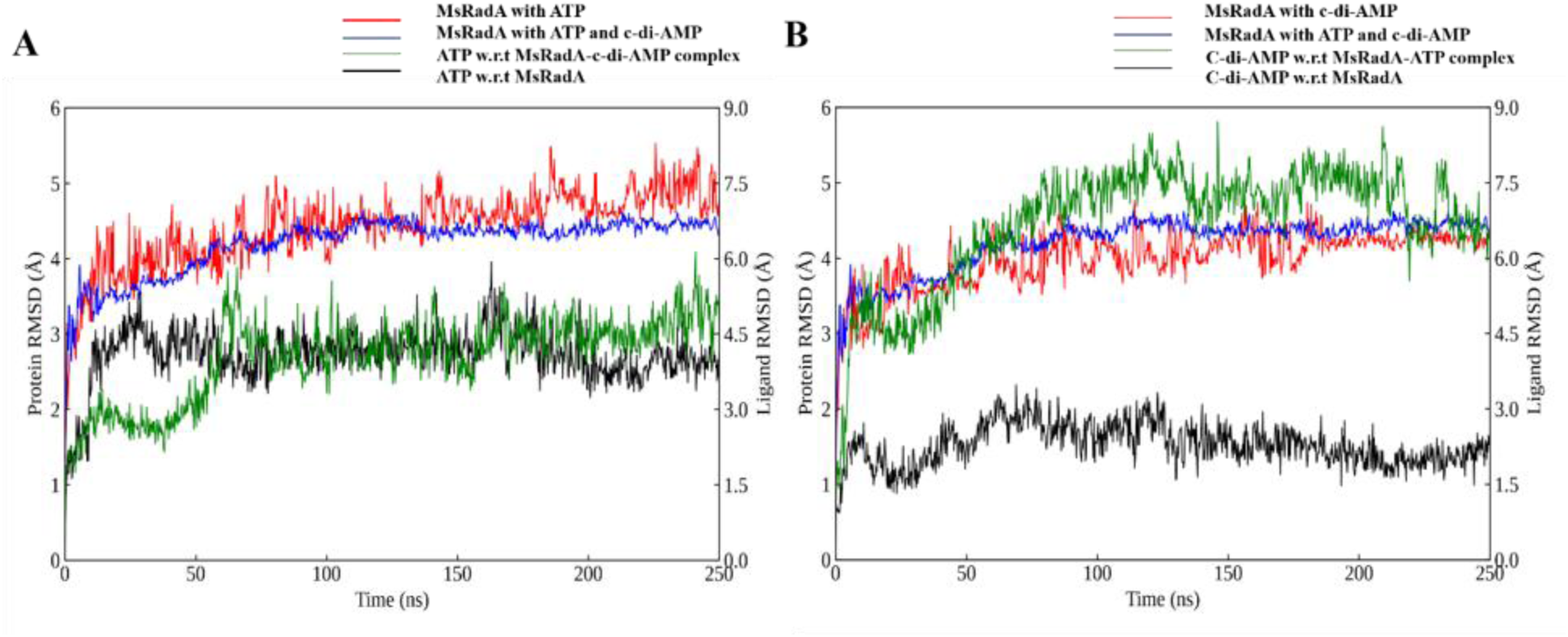
MD simulations of the binary complexes MsRadA-ATP and MsRadA-c-di-AMP complex, and the ternary complex MsRadA-ATP-c-di-AMP. (A) RMSD profiles for ATP-bound MsRadA in the absence and presence of c-di-AMP. Protein RMSD (left axis) is shown for MsRadA with ATP (red) and MsRadA with ATP and c-di-AMP (blue). Ligand RMSD (right axis) is shown for ATP in the MsRadA–c-di-AMP-ATP ternary complex (green) and ATP in the MsRadA-ATP binary complex alone (black). (B) RMSD profiles for c-di-AMP-bound MsRadA in the absence and presence of ATP. Protein RMSD (left axis) is shown for MsRadA with c-di-AMP (red) and MsRadA with ATP and c-di-AMP (blue). Ligand RMSD (right axis) is shown for c-di-AMP in the MsRadA–ATP-c-di-AMP ternary complex (green) and c-di-AMP in the MsRadA-c-di-AMP binary complex (black). Images were produced with OriginPro 9.0.0 (Academic license).

## Discussion

Bacteria rely on several secondary messengers, such as c-di-AMP, to couple cell physiology with the DNA damage response [4], but how this second messenger acts directly on recombination proteins has remained unclear. Based on previous genetic evidence linking RadA to c-di-AMP signaling [14,18], we set out to determine whether MsRadA is a direct binding partner of c-di-AMP, where this interaction occurs relative to the ATP-binding site, and what functional and structural consequences it has. Our ITC data show that c-di-AMP binds MsRadA directly with micromolar affinity (Fig. 2B). This direct interaction with c-di-AMP places RadA alongside RecA as a direct physical target of c-di-AMP within the homologous recombination pathway, rather than a downstream effector regulated solely through RecA. Our finding that RadA independently and directly binds c-di-AMP is consistent with, and mechanistically strengthens, the model that c-di-AMP can directly modulate RadA in the HR machinery, potentially providing coordinated, multi-point control over recombination activity during periods of c-di-AMP accumulation following DNA damage.

A key question was whether c-di-AMP acts through the canonical ATP-binding site or through an independent site. AMP-PCP binding showed the expected strong Mg²⁺-dependence of an ATPase active site (Fig. 1B & 1C), whereas c-di-AMP binding was essentially unaffected by Mg²⁺ (Fig. 2A and Supplementary Fig. S1), arguing against c-di-AMP occupying the same Mg²⁺-coordinated pocket. Similarly, RadA-AMP-PCP complex did not reduce subsequent c-di-AMP binding; the affinity and thermodynamic signature of c-di-AMP binding to the RadA-AMP-PCP complex were nearly identical to those observed for apo RadA, indicating that the two ligands do not compete for occupancy (Fig. 2B). Despite occupying a site distinct from ATP, c-di-AMP reduced the DNA-stimulated ATPase activity of MsRadA at a fixed concentration of c-di-AMP (Fig. 3). Structural modelling of the RadA hexamer placed ATP and c-di-AMP at spatially distinct locations on each subunit interface across all six interfaces, with ATP engaging the Walker A P-loop and an adjacent arginine finger, while c-di-AMP occupied a different position on the same interface (Supplemenatry Fig. S2).

Structural modelling showed that c-di-AMP and DNA are each accommodated at the central channel of the RadA hexamer. The residue-level intermolecular interaction analysis showing the same Arginine and lysine residues contact both c-di-AMP and DNA, engaging the phosphate groups of c-di-AMP and the phosphate backbone of DNA, respectively (Fig. 4A & 4B). This overlap indicates that c-di-AMP and DNA engage a shared structural interface within the central channel of the RadA hexamer, but when DNA is modelled together with c-di-AMP, DNA retains the central channel, and c-di-AMP is instead predicted at a peripheral site proximal to, but distinct from, the ATP pocket (Fig. 5). These models indicate that c-di-AMP does not directly compete with DNA for the same site; rather, DNA occupancy of the central channel is associated with a shift in c-di-AMP’s predicted binding location from the central channel to a peripheral site proximal to the ATP pocket. This DNA-dependent relocation offers a structural model in which c-di-AMP binds a site distinct from the ATP and DNA binding pocket and inhibits DNA-stimulated ATPase, consistent with allosteric modulation of catalysis or of DNA-coupled conformational states rather than of ATP binding. MsRadA might have two c-di-AMP binding sites: one that overlaps the DNA-binding site (Fig. 4) and another that is peripheral to the ATP-binding site (Fig. 5).

This structural independence is consistent with our molecular dynamics data. ATP positioning at the Walker A site was not substantially altered by c-di-AMP, ATP RMSD was ∼4–5 Å both in the presence and absence of c-di-AMP (Fig. 6A), indicating that c-di-AMP does not displace ATP over the 250 ns simulation. In contrast, c-di-AMP showed a markedly higher RMSD in the ternary RadA–ATP–c-di-AMP complex (∼6–8 Å) than when bound to RadA alone (∼2–3 Å) (Fig. 5B). Rather than reflecting competition at a shared site, this pattern is more consistent with ATP occupancy increasing the conformational flexibility of c-di-AMP at the subunit interface of the hexamer, so the stability of bound c-di-AMP depends on the nucleotide state of RadA. Further binding and structural studies are needed to draw firm conclusions about how ATP and c-di-AMP influence each other. The structural and dynamic components of this model rely on predicted complexes and simulation-based analyses rather than experimentally determined structures and should be regarded as a mechanistic hypothesis.

Determining the experimental structures (via Cryo-EM or X-ray crystallography) of MsRadA complexes specifically apo-MsRadA, MsRadA-c-di-AMP, and the ternary MsRadA-ATP-c-di-AMP complex could translate current mechanistic hypotheses into experimentally validated frameworks. Further in *vitro* recombination assays are needed to evaluate how c-di-AMP accumulation directly impacts MsRadA’s core homologous recombination (HR) functions, including DNA strand exchange, D-loop formation, and branch migration. Further characterising the kinetics of c-di-AMP’s inhibition of MsRadA’s ATPase activity under varying DNA lengths and topologies (ssDNA vs dsDNA) would reveal the mechanistic basis of ATPase activity inhibition by c-di-AMP and the response of various DNA structures to c-di-AMP-dependent inhibition kinetics.

## Conclusion

This study identifies MsRadA as a direct, physical binding partner of c-di-AMP, extending the secondary messenger’s regulatory reach within the homologous recombination pathway. C-di-AMP binds MsRadA with micromolar affinity, independently of Mg²⁺, and prior occupancy of the nucleotide site by AMP-PCP does not reduce this binding, indicating that c-di-AMP does not require the ATP-binding pocket. C-di-AMP also partially reduced the DNA-stimulated ATPase activity of MsRadA. Biophysical, functional, and computational data converge on a model in which c-di-AMP occupies a site distinct from the ATP-binding site, and this binding is associated with reduced ATPase activity and potential interference with the DNA-engaged, helical conformation of the RadA hexamer. Together, these data are consistent with a model in which c-di-AMP regulates RadA through a site separate from the active site, either by interfering with DNA engagement or by modulating DNA-coupled conformational states. Whether this occurs allosterically or through competition with DNA remains to be tested. The structural component of this model is based on modelled structures rather than experimentally determined complexes and should be treated as a mechanistic hypothesis.

## Materials and Methods

### Protein Expression and Purification

The *Mycobacterium smegmatis radA* gene (UniProt accession – A0R563) was commercially synthesized and cloned into the pGEX-6P-1 vector by Biotech Desk, generating an N-terminal GST-MsRadA fusion construct. The sequence of the insert was verified by the supplier. The recombinant plasmid was transformed into *Escherichia coli* Rosetta2 (DE3) competent cells by heat shock. Transformants were grown overnight at 37 °C on a petri plate containing Luria-Bertani (LB)-agar supplemented with Ampicillin (100 μg/mL) and chloramphenicol (35 μg/mL). These antibiotics were maintained throughout culturing. Starter culture was used to inoculate four 2 L Erlenmeyer flasks, each containing 1 L of LB broth. These secondary cultures were grown at 37 °C with shaking at 180 rpm until an optical density at 600 nm of approximately 0.8 was reached. Protein expression was induced by the addition of 0.5 mM isopropyl β-D-1-thiogalactopyranoside (IPTG), and the cultures were incubated at 18 °C for 12–15 h.

Cells were harvested by centrifugation and resuspended in lysis buffer containing 50 mM Tris-HCl (pH 8.0), 300 mM NaCl, 1 mM dithiothreitol (DTT), and 5% (v/v) glycerol. The cell suspension was lysed by sonication at 40% amplitude, with 10 seconds on and 20 seconds off in ice and clarified by centrifugation at 14000 rpm at 4 °C for 60 minutes in Sorvall Lynx 4000 Centrifuge (Thermo Fisher Scientific). The supernatant was loaded onto a GST affinity column pre-equilibrated with the lysis buffer. After washing, the bound GST-MsRadA fusion protein was eluted using equilibration buffer supplemented with 10 mM reduced glutathione.

The N-terminal GST tag was removed by incubation with PreScission protease at 4 °C overnight. The cleaved protein was subsequently purified by size-exclusion chromatography using a HiLoad Superdex 200 pg 26/600 column (Supplementary Fig. S3). Fractions corresponding to RadA were pooled and concentrated to approximately 10 mg/ml. Protein concentration was determined by measuring the absorbance at 280 nm using a spectrophotometer, and the purity of the final preparation was assessed by SDS-PAGE (Supplementary Fig. S4)

### Calorimetric Studies

Isothermal titration calorimetry (ITC) experiments were performed at 15 °C using an Affinity ITC LV instrument (TA Instruments, USA). Purified protein (∼50 µM) was loaded into the sample cell, while respective ligands (0.5 – 3 mM) were loaded into the syringe. The reference cell was filled with Milli-Q water.

The sample cell contained 350 μL of either purified MsRadA or the preformed MsRadA-AMP-PCP complex, depending on the experiment. Each titration consisted of 20 sequential injections, comprising an initial 0.5 μL injection followed by 19 injections of 2 μL each, with an interval of 120–150s between injections while stirring at 125 rpm. Appropriate control titrations, including ligand into buffer and buffer into protein, were performed to account for heats of dilution and mixing.

Raw thermograms were acquired using ITCRun software (TA Instruments). Data analysis was performed using Nano Analyze software (TA Instruments) by fitting the integrated heat data to a ‘single set of independent sites’ binding model using nonlinear least-squares regression. The dissociation constant (K_D_), enthalpy change (ΔH) and entropy change (ΔS) were treated as adjustable parameters during curve fitting.[24–26]

### ATPase activity assay

ATPase activity was measured using an NADH-coupled spectrophotometric assay in 96-well plates (150 µL reaction volume) at 37 °C, using a multimode plate reader. Reaction mixtures contained 0.5 µM protein, 2.5 µM single stranded DNA (45-mer) (Supplementary Table 2), 10 mM Tris–Cl (pH 7.5), 5 mM Mg(OAc)₂, 0.1 mM DTT, 3 mM phosphoenolpyruvate (PEP), 0.2 mg mL⁻¹ NADH, 20 U mL⁻¹ Pyruvate kinase, 10 U mL⁻¹ L-lactate dehydrogenase and 2 mM ATP, coupling ATP hydrolysis to NADH oxidation. The decrease in absorbance at 340 nm was monitored continuously.

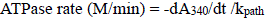

where k_path_ is the molar absorption coefficient of NADH corrected for the optical path length of the well; for a 150 µL fill volume, k_path_ = 2,383 M⁻¹ (empirically determined calibration factor obtained from an NADH standard curve).

The effect of c-di-AMP on ATPase activity was assessed by comparing initial velocities of reactions containing fixed concentration of c-di-AMP (0.5 mM in this case) and in the absence of it as a control under identical ATP conditions.

Reaction velocities were calculated from the linear portion of the NADH depletion curves (corresponding to <20% total NADH consumption) using ICEKAT (Initial and Continuous Enzyme Kinetics Analysis Tool). [27–29]

### Structural modeling of the MsRadA hexamer in complex with ligands and DNA

To generate a structural model of the fully liganded MsRadA hexamer, the MsRadA protein sequence was submitted to the Protenix server, an open-source, all-atom biomolecular structure prediction model capable of jointly predicting protein, ligand, and ion complexes in a single inference run. [30] The hexameric MsRadA assembly was modeled in complex with six molecules of ATP, six Mg²⁺ ions, and six molecules of c-di-AMP (one full ligand set per protomer) and a DNA duplex with an overhang (Supplementary Table 2). The resulting models was ranked by the server’s confidence metrics, and the top-ranked prediction was used for subsequent analyses, including identification of binding sites by using PLIP (Protein-Ligand Interaction Profiler) server [31].

### Molecular Dynamics Simulations of MsRadA complexes

The modeled structure of MsRadA complexes (MsRadA-ATP-c-di-AMP ternary complex, MsRadA-ATP and MsRadA-c-di-AMP binary complexes) was imported into Maestro (Schrodinger Suite, Release 2022-1, Academic license) and prepared using the Protein Preparation Wizard, which fixed structural issues and optimized hydrogen bonding. A simulation-ready system was then built using the System Builder tool. Na⁺ and Cl⁻ ions were added to neutralize the overall charge, and the complex was placed in an orthorhombic box with a 10 Å buffer on all sides. The system was solvated with TIP3P water and 150 mM NaCl to mimic physiological salt conditions. Energy minimization was carried out using the OPLS4 force field. Prior to production simulations, each solvated system was relaxed using a stepwise equilibration protocol in which positional restraints on the solute were progressively removed, allowing the solvent and ions to adapt to the solute before the protein–ligand complex was allowed to move freely. This reduces steric clashes and prevents structural distortion from the abrupt release of a system that has not yet been thermally and volumetrically equilibrated. The production simulation ran for 250 ns under NPT conditions (300 K, 1 atm), using a Nose-Hoover thermostat and Martyna-Tobias-Klein barostat to maintain temperature and pressure. Long-range electrostatics were handled with the Particle Mesh Ewald (PME) method (9 Å cutoff), and a 2 fs timestep was used with the RESPA integrator. Trajectory analysis and visualization of MsRadA’s interactions with ATP and c-di-AMP were performed using Maestro and PyMOL (Version 2.5, Schrödinger, LLC). [32–34]

### Author contributions

ER contributed to the conceptualization, funding acquisition, and supervision, writing, review and editing. SS Investigation, Data curation, Formal analysis, Visualization, Writing-review & editing. SPG contributed data curation, formal analysis, writing-review, and editing. KN Formal analysis, writing-review, and editing.

## Acknowledgments

SS would like to thank the University Grants Commission (UGC), Government of India, for the fellowship. SPG would like to thank the Department of Biotechnology (DBT), Government of India, for the fellowship.

## Data availability statement

The data that support the findings of this study are available upon request from the corresponding author.

## Conflict of Interest

The authors declare that there are no conflicts of interest.

## Abbreviations

Ms: *Mycobacterium smegmatis*
ITC: Isothermal Titration Calorimetry
AMP-PCP: β,γ-methylene adenosine 5′-triphosphate
MD: Molecular Dynamics
HR: Homologous Recombination
ATP: Adenosine Triphosphate
c-di-AMP: Cyclic di adenosinediphosphate
c-di-GMP: Cyclic di guanosinediphosphate
CDN: Cyclic di nucleotide
DisA: DNA integrity Scanning A
RadA: Radiation sensitive A
RecA: Recombinase A
K_D_: Molar dissociation constant
RMSD: Root mean square deviation
DNA: Deoxyribonucleic acid
pTM: Predicted TM-score
ipTM: Interface predicted TM-score

## Supporting Information

**Supplementary Figure S1.**
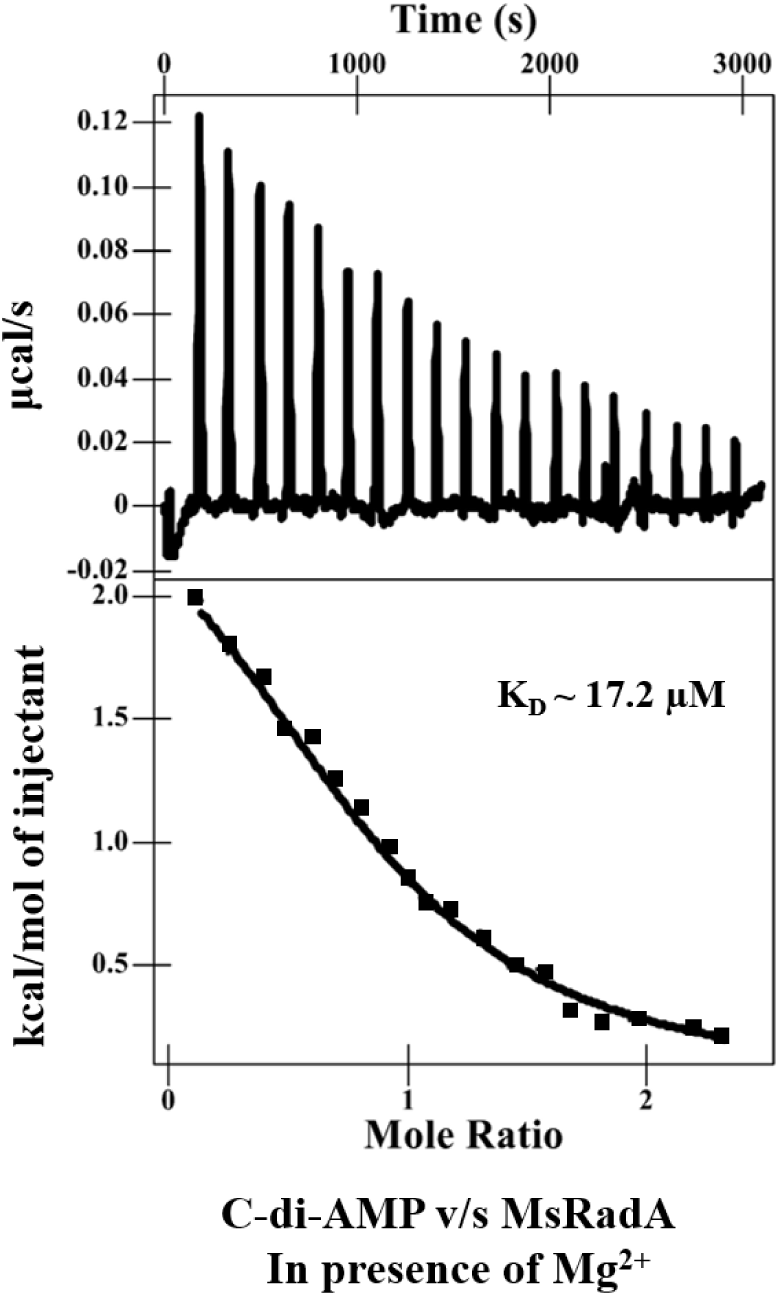
ITC binding study of C-di-AMP against the MsRadA in the presence of magnesium.

**Supplementary Figure S2.**
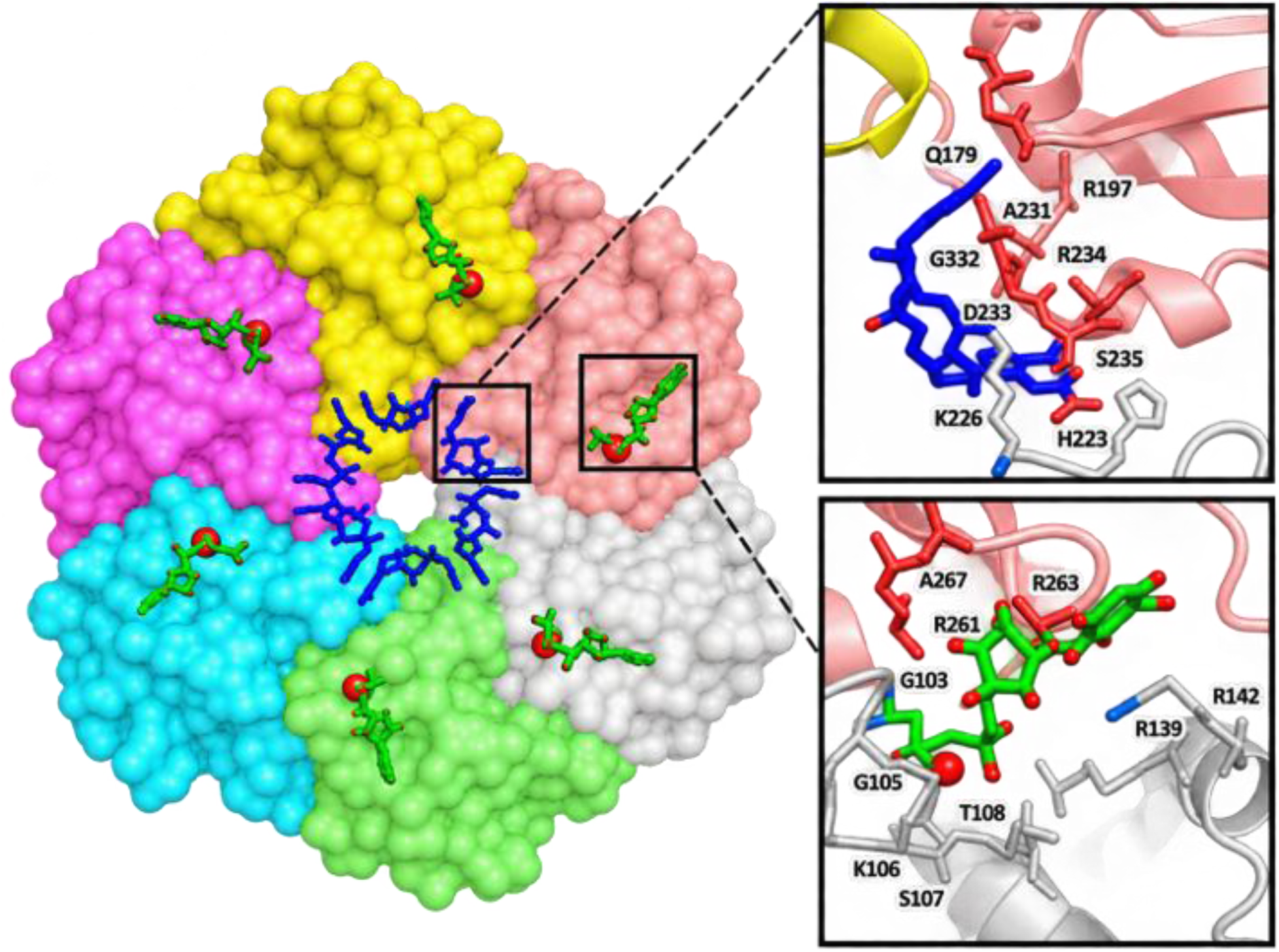
Structural modeling of the MsRadA hexamer bound c-di-AMP and ATP. MsRadA hexameric ring (subunits A-F, individually colored), green sticks at each subunit interface represent ATP (green sticks) occupying the canonical nucleotide-binding pocket and c-di-AMP (blue sticks) at the center of the channel.

**Supplementary Figure S3.**
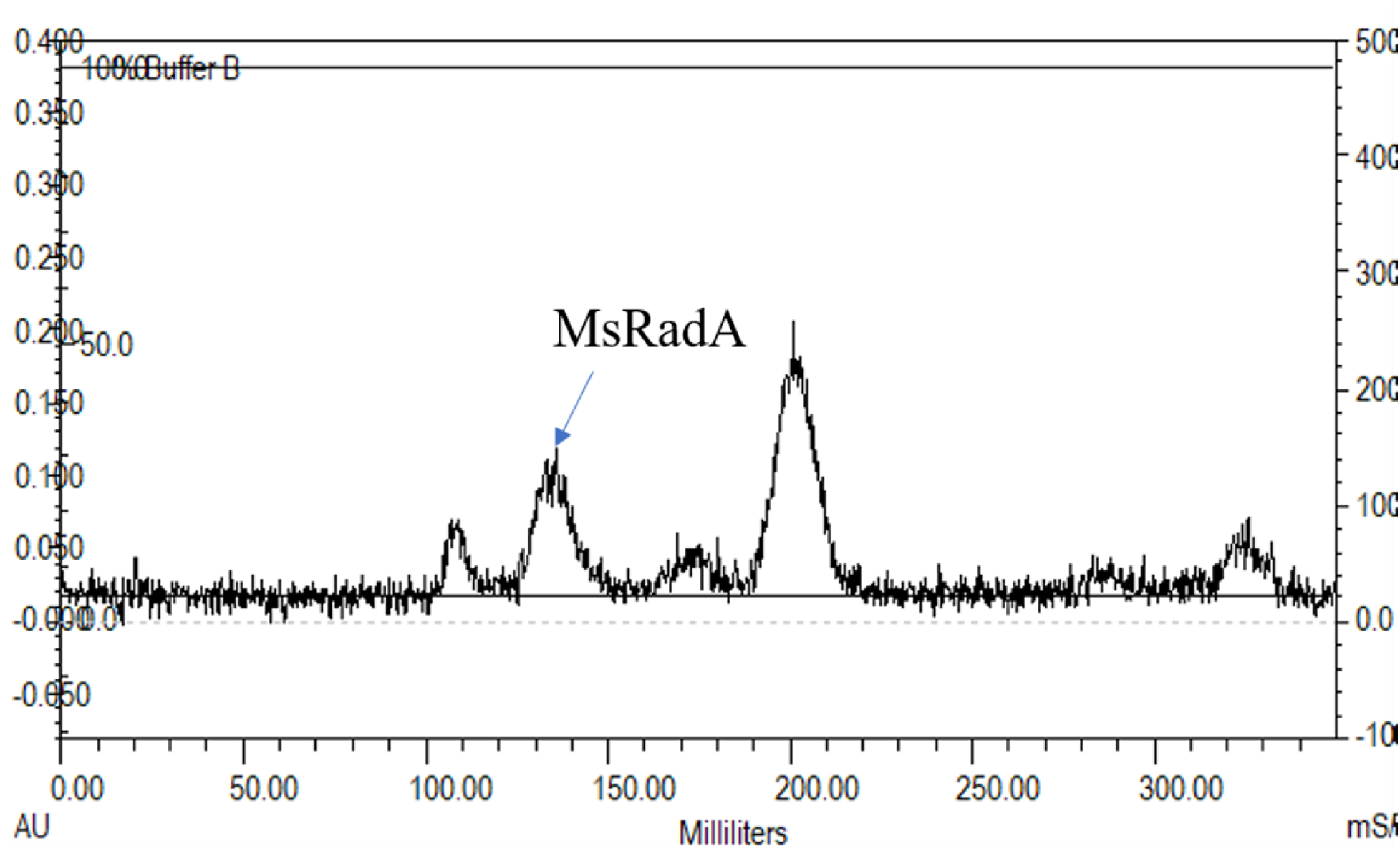
Purification of MsRadA. Size Exclusion Chromatogram (SEC) of MsRadA. The elution profile (absorbance at 280 nm, AU, versus elution volume in mL) shows the purification of *Mycobacterium smegmatis* RadA (MsRadA) after GST-tag cleavage. The peak at ∼130 mL corresponds to MsRadA (arrow). The peak to its left, eluting at ∼105 mL near the void volume, represents aggregated protein. The peak to its right, at ∼200 mL, corresponds to the cleaved GST tag. The concentrated sample was loaded automatically onto the SEC column from the injector loop by BioRad DuoFlo pump.

**Supplementary Figure S4.**
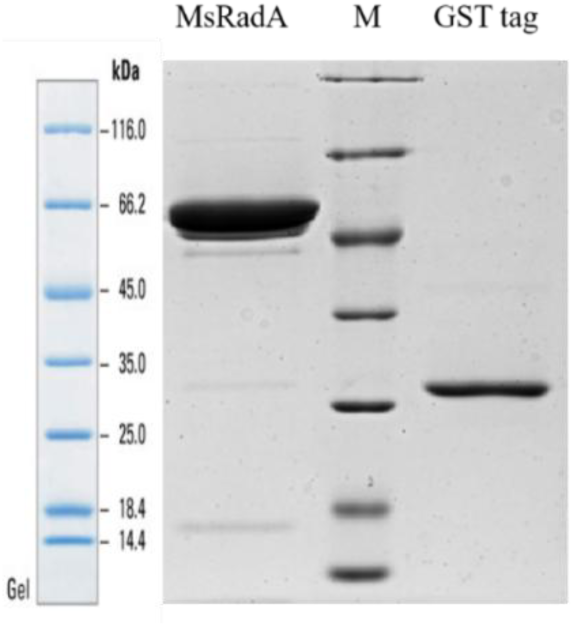
SDS-PAGE analyses purified *Ms*RadA. Lane 1-MsRadA (49kDa), Lane 2-Protein Marker and Lane 3-GST tag after cleavage.

**Supplementary Table 1.** Summary of binding thermodynamics determined with ITC.

| Titration at 288K |  | Thermodynamics |  |  |  |  |
| --- | --- | --- | --- | --- | --- | --- |
| Syringe | Cell | $\Delta H$ (kcal mol<sup>-1</sup>) | $\Delta S$ (cal mol<sup>-1</sup> K<sup>-1</sup>) | $-T\Delta S$ (kcal mol<sup>-1</sup>) | $\Delta G$ (kcal mol<sup>-1</sup>) | $K_D$ (μM) |
| AMP-PCP | MsRadA | -1.10 | 14.7 | -4.24 | -5.34 | 88.9 |
| AMP-PCP+Mg<sup>2+</sup> | MsRadA + Mg<sup>2+</sup> | +0.81 | 26.5 | -7.64 | -6.84 | 6.52 |
| C-di-AMP | MsRadA | +2.15 | 29.6 | -8.54 | -6.39 | 14.2 |
| C-di-AMP+Mg<sup>2+</sup> | MsRadA + Mg<sup>2+</sup> | +3.75 | 35.1 | -10.12 | -6.37 | 14.7 |
| C-di-AMP+Mg<sup>2+</sup> | MsRadA + AMP-PCP+Mg<sup>2+</sup> | +2.83 | 31.6 | -9.11 | -6.28 | 17.2 |

**Supplementary Table 2.**
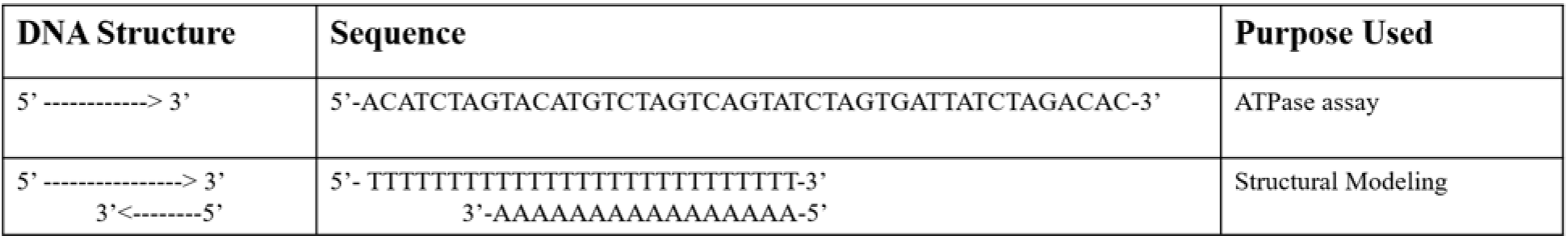
List of DNAs used in our study.

**Supplementary Table 3.** Table of the confidence scores of the 5 models. The pTM and ipTM scores of each model are mentioned.

| **S no.** | **Model** | **pTM** | **ipTM** |
| --- | --- | --- | --- |
| 1. | MsRadA-ATP | 0.80 | 0.78 |
| 2. | MsRadA-c-di-AMP | 0.70 | 0.66 |
| 3. | MsRadA-ATP-c-di-AMP | 0.77 | 0.75 |
| 4. | MsRadA-ATP-DNA | 0.76 | 0.74 |
| 5. | MsRadA-ATP-DNA-c-di-AMP | 0.74 | 0.71 |

## Notes

### Competing Interest Statement

The authors have declared no competing interest.

